# Gene regulatory network analysis of somatic embryogenesis identifies morphogenic genes that increase maize transformation frequency

**DOI:** 10.1101/2025.06.22.659756

**Authors:** Jim Renema, Svitlana Lukicheva, Isabelle Verwaerde, Stijn Aesaert, Griet Coussens, Jolien De Block, Carolin Grones, Thomas Eekhout, Bert De Rybel, Rhoda A.T. Brew-Appiah, Christopher A. Bagley, Lennart Hoengenaert, Klaas Vandepoele, Laurens Pauwels

## Abstract

Somatic embryogenesis allows a somatic plant cell to develop into an embryo, and potentially into a fertile plant. Transcription factors such as BABY BOOM can induce somatic embryogenesis when ectopically expressed and are widely used for aiding regeneration in tissue culture for transformation and gene editing of crops. Nevertheless, regeneration remains a bottleneck and alternative morphogenic genes are highly desired. Here, we co-expressed BABY BOOM and WUSCHEL2 in zygotic maize (*Zea may*s L.) embryos and studied gene regulatory networks in induced somatic embryos at the single-cell level. By inferring cell-type-specific regulons, we prioritized candidate regulators and confirmed functionality of four transcription factors, bHLH48, EREB152, GRF4, and HB77, for enhanced maize transformation frequency, leading to fertile, transgenic plants. Interestingly, the basic helix-loop-helix and homeodomain-leucine zipper families had previously not been associated with induced somatic embryogenesis. Our work will contribute to more efficient transformation, much needed to deliver on the promise of gene editing for agriculture.

## Introduction

Gene editing enables the translation of basic research discoveries into crop varieties tailored to meet agricultural demands. For cereal crops like *Zea mays* (maize), immature zygotic embryos are used as explants and co-cultured with *Agrobacterium tumefaciens* to introduce a T-DNA encoding CRISPR/Cas9-components to cells. Using tissue culture, cells are grown as undifferentiated callus and transgenic plant cells selected using herbicides or antibiotics. Addition of synthetic auxins induces the formation of somatic embryos (SEs) that may germinate to transgenic plantlets (Kaush *et al*., 2021; Lee and Wang, 2023). Low transformation frequencies (the fraction of explants generating one or more transgenic plants) have hampered use of maize in research and industry. Moreover, only a fraction of transgenic events are of high quality, containing only a single T-DNA and no vector backbone (Vandeputte *et al*., 2024).

Efforts for improvement include equipping *Agrobacterium* strains with ternary vector systems to increased virulence (Zhang *et al*., 2019, Vandeputte *et al*., 2024, Aliu *et al*., 2024) and promoting direct somatic embryogenesis through the co-expression of transcription factors (TF) known as morphogenic regulators (MRs). A breakthrough in maize was the expression of the maize APETALA2/ ETHYLENE RESPONSIVE ELEMENT BINDING PROTEIN (AP2/EREBP) factor BABY BOOM (ZmBBM) and the maize homeobox factor WUSCHEL 2 (ZmWUS2) as additional genes on the T-DNA (Lowe *et al*., 2016). Initially, *ZmBBM* and *ZmWUS2* were respectively expressed using the constitutive maize ubiquitin (ZmUBI) and the *Agrobacterium* nopaline synthase (nos) promoters (Lowe *et al*., 2016). This increased both transformation frequency and gave more flexibility to genotypes that could be used but were often paired with pleiotropic developmental effects including infertility (Lowe *et al*., 2016, Lowe *et al*., 2018). Hence, MRs were flanked by loxP sites and combined with a Cre recombinase, driven by an inducible promoter. This enabled the auto-excision of the MRs after initiation of somatic embryogenesis. Secondly, *ZmBBM* was expressed using the PHOSPHOLIPID TRANSFER PROTEIN (ZmPLTP) promoter, driving expression in the scutellum epithelium of the embryo explant, and *ZmWUS2* was driven by the auxin-inducible promoter of *ZmIAA25* (pZmAXIG1). Transformation of pZmPLTP::ZmBBM pZmAXIG::ZmWUS2 prompted the formation of SEs within a few days (Lowe *et al*., 2018). Other MRs have been described to aid transformation frequency such as ZmWOX2A, a WUS-related protein (McFarland *et al*., 2023), and the *ZmBBM*-related gene *ZmBBM2* (Du *et al*., 2019). A fusion protein between a GROWTH-REGULATING FACTOR (TaGRF4) and GRF-INTERACTING FACTOR (TaGIF1) protein was found to improve regeneration in wheat (Debernardi *et al*., 2020) and could be translated to maize (Vandeputte *et al*., 2024). Finally, a transcriptional regulatory network analysis of wheat regeneration identified two DOF-factors (Liu *et al*., 2023). These and other studies suggest the potential for discovering additional TF that could serve as valuable biotechnological tools (Jiang *et al*., 2024).

We here leveraged the ability of ZmBBM and ZmWUS2 to induce SEs to identify maize basic helix-loop-helix 48 (bHLH48), EREB152, GRF4, and HD-Zip 77 (HB77) as MRs that improve maize transformation frequency.

## Results

### Single-cell RNA-sequencing of induced somatic embryogenesis

Expressing pZmPLTP::ZmBBM pZmAXIG1::ZmWUS2 in scutellum epithelium cells leads to the rapid formation of SEs (Lowe *et al*., 2018, Aesaert *et al*., 2022). By combining these genes with the ternary vector pVS1-VIR2, we can drastically increase the number of transformed cells and thus SEs per explant (Fontanet-Manzaneque *et al*., 2024). We constructed the binary plasmid pG3R-WOY-SI that contains, in addition to pZmPLTP::ZmBBM pZmAXIG1::ZmWUS2, two reporters: the *E. coli* β-GLUCURONIDASE (GUS) and the YELLOW FLUORESCENT PROTEIN with nuclear localization signal (YFP-NLS, Fig. 1A). At seven days after transformation (DAT), and even more prominently at 10 DAT, a high number of multicellular SEs can be observed per zygotic embryo explant (Fig. 1B). At 7 DAT, we isolated protoplasts from approximately 300 embryo explants and enriched YFP-NLS-expressing protoplasts using Fluorescent Activated Cell Sorting (FACS). This yielded approximately 100,000 YFP-NLS-expressing protoplasts that were used for 10x Genomics-based single cell partitioning and library preparation (Fig. 1C). Following quality control and filtering steps, we retained 6,830 high-quality cells for downstream analysis. After cell cycle regression, we identified 17 clusters that were visualized using Uniform Manifold Approximation and Projection (UMAP) and annotated with markers from maize literature when possible (Fig. 2A, Extended Data Fig. 1, Supplementary Table 1A). In conclusion, we established an efficient strategy for studying induced somatic embryogenesis at the single-cell level by selectively isolating SE cells based on fluorescence.

**Figure 1.**
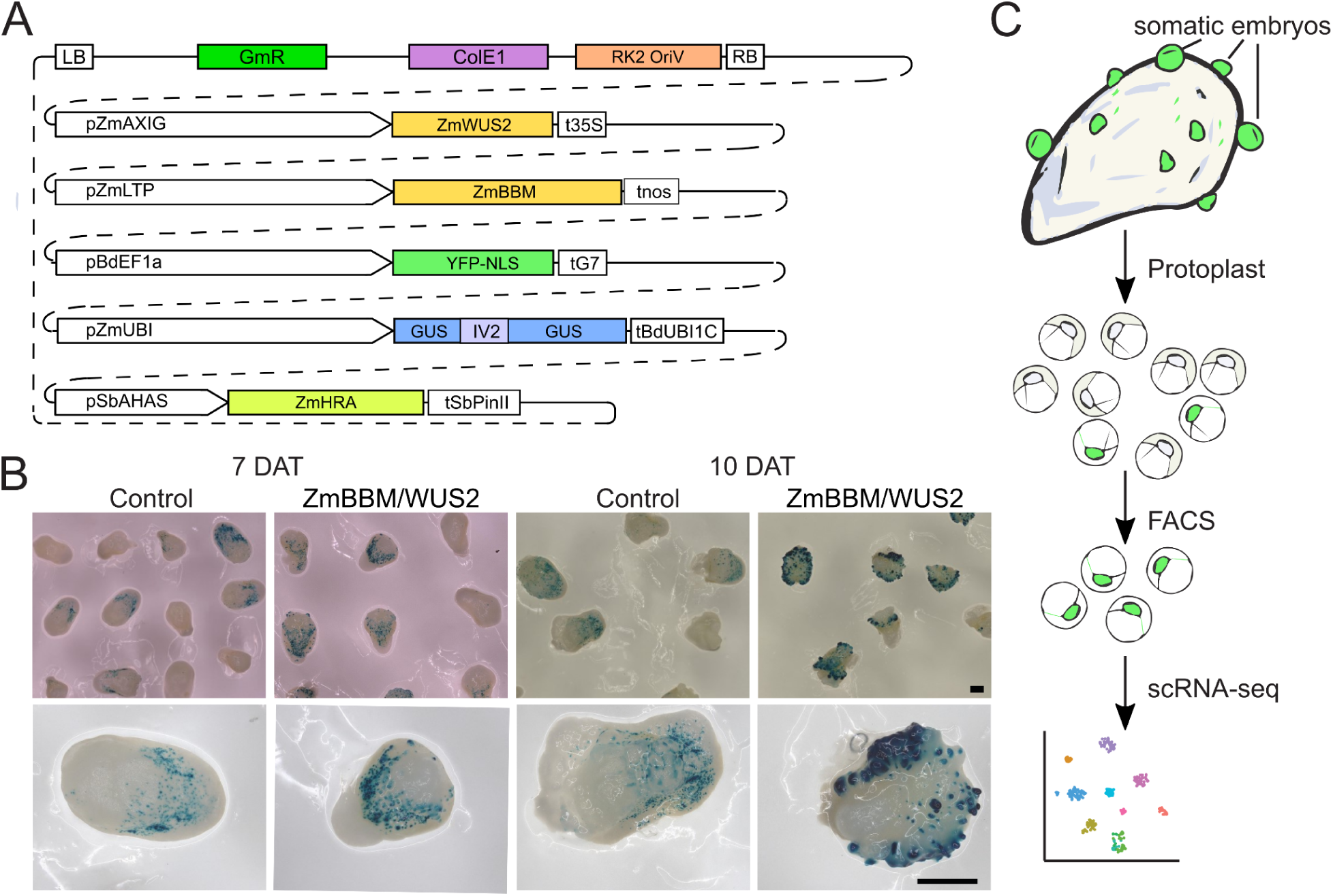
Strategy to generate a single-cell RNA sequencing dataset of induced somatic embryo formation in maize. **A**, Diagram of the binary vector pG3R-WOY-SI. GmR, Gentamicin resistance; ColE1 and RK2 OriV, origin of replication sequences; RB, right border; pZmAXIG, the auxin-inducible promoter of ZmIAA25; ZmWUS2, maize WUSCHEL; t35S, pCaMV 35S terminator; pZmLTP, promoter of the maize PHOSPHOLIPID TRANSFER PROTEIN; ZmBBM, maize BBM; tnos, *Agrobacterium* nopaline synthase terminator; pBdEF1a, *Brachypodium distachyon* ELONGATION FACTOR 1a promoter; YFP-NLS, YELLOW FLUORESCENT PROTEIN fused to a nuclear localization signal; tG7, *Agrobacterium* G7 terminator; pZmUBI, maize UBIQUITIN promoter; GUS, β-GLUCURONIDASE with intron IV2 of the potato gene ST-LS1; tBdUBI1C, *Brachypodium distachyon* UBIQUITIN1 terminator; pSbAHAS, *Sorghum bicolor* ACETOHYDROXYACID SYNTHASE; ZmHRA, maize HIGHLY RESISTANT AHAS; tSbPinII, potato PinII terminator. **B**, representative images of immature maize embryos transformed with pG3R-WOY-SI or a GUS-only control 7 or 10 days after transformation (DAT) and stained for GUS expression. Scale bars: 1 mm. **C**, overview of the experimental setup. Protoplast isolation is performed starting from immature embryos transformed with pG3R-WOY-SI at 7 DAT. YFP-expressing protoplasts are sorted using fluorescence activated cell sorting (FACS) and used for single-cell RNA sequencing.

**Figure 2.**
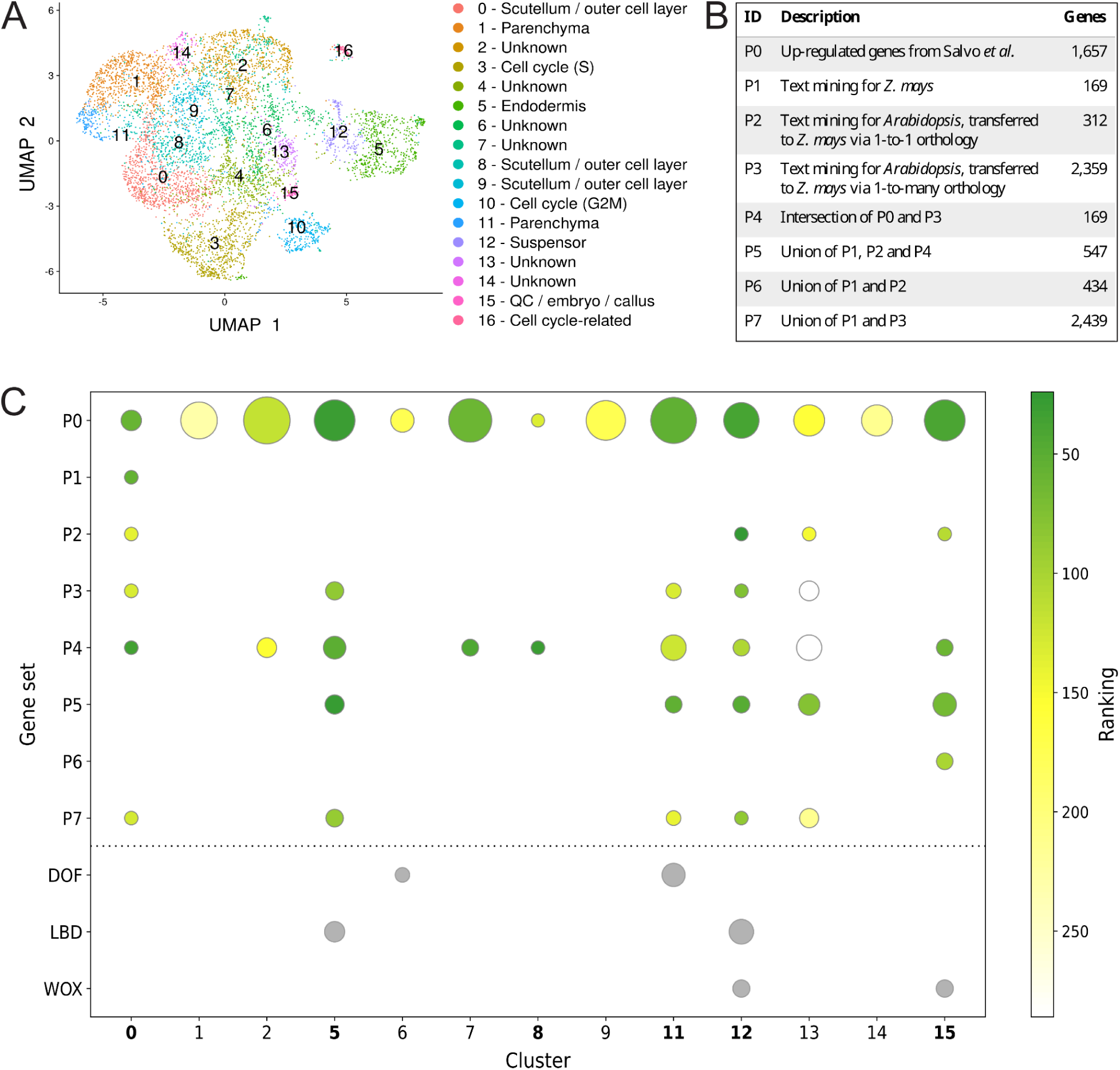
Prioritization of clusters associated with somatic embryogenesis in the scRNA-seq dataset. **A**, UMAP plot illustrating the clusters identified in the scRNA-seq dataset. Each point represents a single cell, colored by cluster identity. **B**, Gene sets (priors) related to somatic embryogenesis, collected from various sources, used for MINI-EX runs. Columns indicate prior IDs, descriptions, and the number of genes in each set. **C**, Single-cell clusters associated with somatic embryogenesis were identified through: 1) enrichment analysis of the top 100 upregulated genes of each cluster against the eight priors (P0-P7) and relevant TF families (DOF, LBD, and WOX); enrichment is visualized as circles, where size reflects the -log10 of the enrichment q-value (larger circles indicate higher enrichment), and 2) MINI-EX median rank of the gold standard TFs associated with somatic embryogenesis assigned using each prior, with greener circles indicating better ranks (TF families were not used as priors and are depicted in gray). Clusters with prominent green circles were identified as associated with somatic embryogenesis and are shown in bold.

### Prioritization of single-cell clusters using text mining

As little information was readily available from the literature to annotate the clusters, we used automated text mining to prioritize clusters most likely involved in somatic embryogenesis. We compiled gene sets known to be associated with somatic embryogenesis and related processes in maize and *Arabidopsis thaliana* (hereafter *Arabidopsis*). Ontology terms, such as ‘somatic embryogenesis’, ‘meristem development’, and ‘regeneration’, were used to extract genes from the literature (Supplementary Table 1B). This information was collected as species-gene-trait triples, (*e.g.*, maize-BBM-somatic embryogenesis). For each triple, we documented publications, the total number of occurrences, and the sections (*e.g.*, abstract) and used these metrics to filter and reduce the number of false-positive triples. Based on manual curation of a subset, we estimate that 81% of the triples are correct.

From the filtered triples, we derived three gene sets, referred to as *priors*, as they represent the current knowledge about somatic embryogenesis. The first prior, termed P1, consisted of 247 triples associated with 169 unique genes identified for maize. The second prior, termed P2, included triples from *Arabidopsis*, with gene identifiers translated to maize using one-to-one gene orthology, and comprised 418 triples associated with 279 unique genes. The third prior, termed P3, consisted of *Arabidopsis* triples translated to maize using one-to-many gene orthology, resulting in 4,017 triples associated with 2,344 unique genes (Fig. 2B). In addition to the three priors collected via text mining, we included differentially expressed genes from a bulk RNA-seq dataset of maize somatic embryogenesis (Salvo *et al*., 2014) as prior P0. Finally, we generated various combinations of the four initial priors, resulting in 8 final priors, and supplemented them with 38 manually verified genes (Supplementary Table 1C) from P1 (Fig. 2B).

To identify the most relevant clusters, we performed an enrichment analysis of the top 100 up-regulated differentially expressed genes (DEGs) in each cluster against the text-mined priors P1-P3. Clusters 0, 5, 11, 12, 13, and 15 were enriched for one or more of the priors (Fig. 2C). Additionally, we performed enrichment against TF families known to be involved in the processes of interest, specifically WUSCHEL-related homeobox (WOX, Wang *et al*., 2022), Lateral Organ Boundaries domain (LBD, Joshi *et al*., 2023) and DNA-binding with One Finger (DOF, Liu *et al*., 2023). Clusters 5, 6, 11, and 15 showed enrichment for at least one of the TF families (Fig. 2C). Taken together, we selected clusters 0, 5, 8, 11, 12, and 15 for further analysis.

### Prioritization of candidate regulators using cell-type-specific gene regulatory networks

To subsequently prioritize candidate regulators within the relevant single-cell clusters, we performed cell-type-specific gene regulatory network inference using MINI-EX (Ferrari *et al*., 2022; Staut *et al*., 2023). MINI-EX first constructs a coexpression network from a single-cell dataset, which is composed of regulons, each representing a TF and its set of target genes. This initial network is further refined to retain only TF motif-supported interactions between TFs and their target genes, based on TF binding site enrichment analysis per regulon. The resulting regulons are assigned to specific clusters through enrichment analysis based on the differential expression of their target genes, and subsequently these cell-type specific regulons are filtered for the TF’s expression within their clusters. The final regulons are ranked based on their network properties, their cluster specificity, and their enrichment for traits of interest in their target gene ontology. Since each cluster identified as related to somatic embryogenesis was enriched for different priors (Fig. 2B), we executed MINI-EX on each prior and evaluated the performance of each run. The performance of the eight priors was evaluated using a gold standard of 36 manually curated TFs from the literature (Supplementary Table 1D). For each MINI-EX run, we evaluated the median ranking of these gold standard TFs for the clusters of interest. As expected, the best performing priors differed per cluster: for example the best TFs ranking for clusters 0 and 8 was observed with priors P1 and P4; while for clusters 5 and 12, with priors P2 and P5 (Fig. 2C, Supplementary Table 1E). This result indicates that the prioritization of candidate TFs should be done using different priors for each cluster. Candidate master regulators were selected within the top 50 for all the associated priors, and we included limited gene size and complexity for gene synthesis as additional technical criteria. Finally, we concluded on 60 TFs for experimental follow-up (Supplementary Table 1F).

### Rapid and Visual Screening of Somatic Embryogenesis Using the RUBY Reporter

The RUBY reporter (He *et al*., 2020) has been used successfully to evaluate maize transformation (Lee *et al*., 2023). Expression using the 35S promoter was however associated with reduced plant growth (Lee *et al*., 2023). Here, we used the weaker constitutive switchgrass (*Panicum virgatum*) ubiquitin promoter (PvUBI2, Mann *et al*., 2011) to drive RUBY and generate the binary destination vector pG3B-SI-AG-RUBY. When combined with (candidate) MRs, SE formation can now be monitored in a non-destructive manner over time using light microscopy (Fig. 3A). For validation, we expressed ZmBBM, ZmWUS, ZmBBM fused to ZmWUS2 by a P2A peptide (Zhang *et al*., 2019) or tdTomato (tdT) under control of the ZmUBI promoter (Fig. 3A-B). In addition, we co-expressed ZmBBM with a ZmUBI promoter and ZmWUS2 with a nopaline synthase promoter (pnos), further referred to as ZmBBM/WUS2. Starting eight days after transformation, we observed the emergence of embryo-like structures on the scutellum epithelium (Fig. 3B). For all constructs containing ZmBBM, the structures accumulated high amounts of betalains. When expressing ZmWUS2 alone, morphogenic structures were often negative for RUBY expression. We attribute this to the well-known ability of ZmWUS2 to stimulate meristem formation in a non-cell autonomous manner (Hoerster *et al*., 2020). Hence, pPvUBI2::RUBY is an excellent marker to monitor early events during MR-induced somatic embryogenesis in maize.

**Figure 3.**
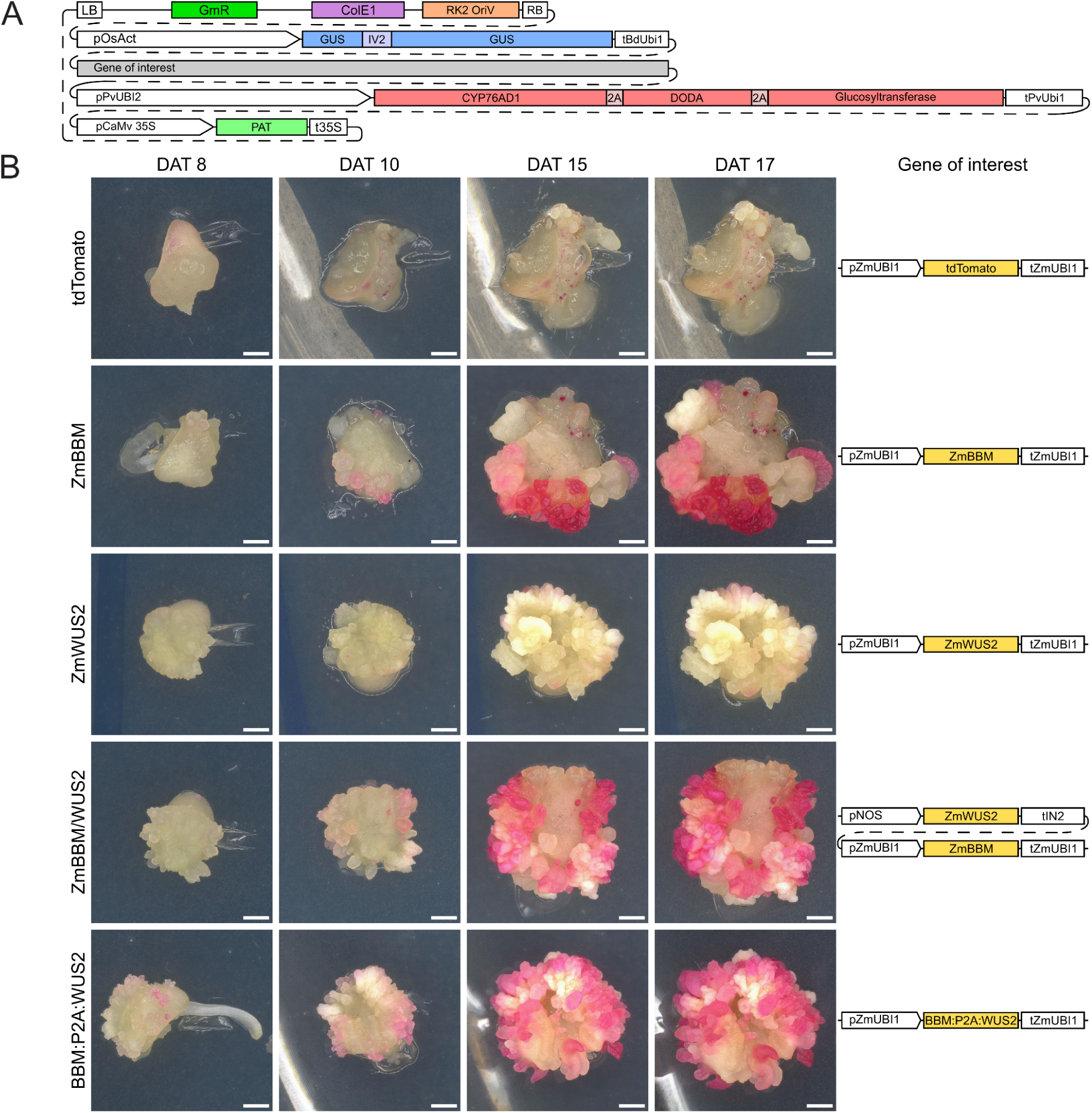
RUBY as a marker to monitor morphogene-induced somatic embryogenesis in maize. **A**, Diagram of the pG3B-SI-AG-RUBY reporter with genetic elements as in Figure 1 in addition to pPvUBI2 and tPvUBI1, *Panicum virgatum L.* UBIQUITIN promoter and terminator; CYP76AD1, P450 CYP76AD1; 2A peptide; DODA, l-DOPA 4,5-dioxygenase; pCaMv 35S and t35S, CaMV 35S promoter and terminator; PAT, PHOSPHINOTRICIN ACYLTRANSFERASE. The Gene of interest is a cloning site consisting of the superfolder GFP (sfGFP) expressed by the *Escherichia coli* glpT promoter and flanked by Green Gate A and G overhangs. **B**, B104 immature embryos were transformed using a construct containing both the non-destructive RUBY reporter and a morphogenic regulator cassette as shown in “Gene of interest”. Shown are representative embryos followed over time using digital light microscopy. The control *tandem dimer Tomato* (*tdTomato*), maize *BABY BOOM* (*ZmBBM*), maize *WUSCHEL2* (*ZmWUS2*), and BBM*:P2A:WUS2* are expressed under control of the maize *UBIQUITIN1* promoter (pZmUB1I) and terminator (tZmUBI1). The representative images of ZmBBM/WUS2 construct contains ZmBBM under control of the maize *UBIQUITIN1* promoter (pZmUB1I) and terminator (tZmUBI1) in addition to *ZmWUS2* under the control of the *Agrobacterium tumefaciens* nopaline synthase promoter (pnos) and maize In2-1 terminator. DAT, days after transformation. Scale bars: 1 mm.

For 60 candidate TFs (Supplementary Table 1F) the coding sequences were domesticated, synthesized, and cloned under control of ZmUBI promoter and terminator in the binary vector pG3B-SI-AG-RUBY (Supplementary Table 2, Fig. 3A). In a primary screen of the 60 candidates, most candidates produced phenotypes comparable to the tdT control. However, several candidates displayed clear morphological changes. Expression of three candidates, *bHLH121* (Zm00001eb102260), *bHLH136* (Zm00001eb317460), and *bHLH29* (Zm00001eb252100), was associated with the appearance of morphogenic structures expressing RUBY at 10 DAT (Extended Data Fig. 2A). Notably, all three belong to the same basic helix-loop-helix (bHLH) TF subfamily (Extended Data Fig. 2B). Nevertheless, these structures did not develop further and we discarded these candidates for our purpose. Remarkably, we obtained several candidates which displayed obvious ZmBBM/WUS2-like phenotypes with the formation of embryo-like structures (Fig. 4, Extended Data Fig. 3). These TFs were not specifically associated with a specific cluster, and each of the selected clusters associated with at least one candidate (Supplementary Table 1F). To quantify the responses, we determined the number of zygotic embryos developing RUBY-expressing structures at 13 DAT. While ZmBBM/WUS2 induced morphogenic structures in 91% of embryos, the tdT negative control averaged 8%. Among the candidates, response was significantly increased ranging from 33% for GRF5 (Zm00001eb430300), to 77% for HB77 (Zm00001eb377480), and 91% for EREB152 (Zm00001eb389400) (Fig. 4, Supplementary Table 3). Based on the primary results, twenty candidates were selected for a confirmation screen. This confirmed a reproducible and significant induction for bHLH48 (Zm00001eb062130), EREB152, GRF4 (Zm00001eb052070), GRF5, and HB77 (Supplementary Table 4).

**Figure 4.**
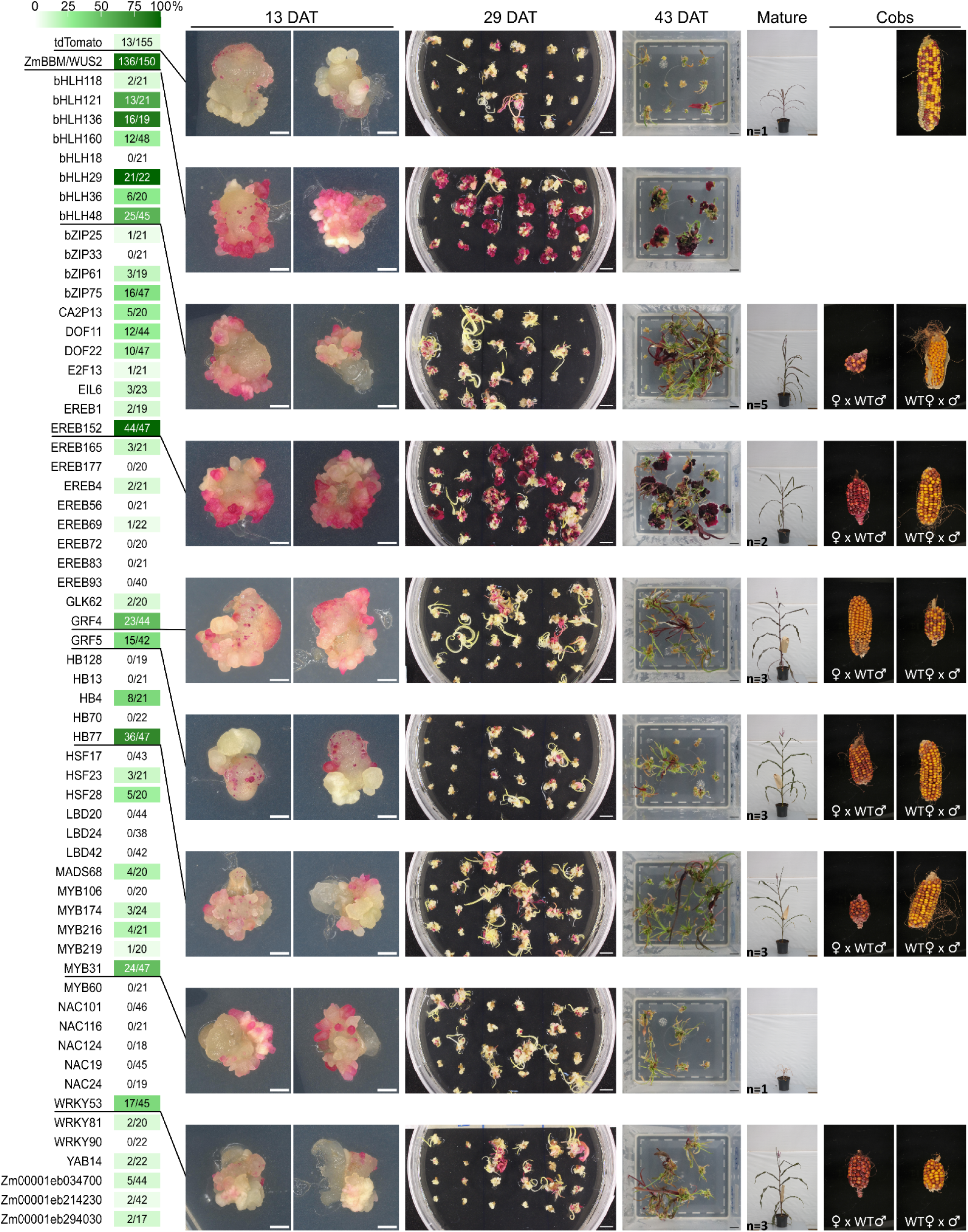
Primary screen using the RUBY reporter identifies candidate morphogenic regulators in maize. B104 immature embryos were transformed using a binary vector containing the pPvUBI2::RUBY::tPvUBI1 reporter and coding sequence of one out of 60 candidate transcription factors under control of the ZmUBI1 promoter and terminator, sorted per protein family. Shown are representative images of the negative tdT control, ZmBBM/WUS2, and 7 candidates that were identified and monitored during the primary screen to have a response similar to ZmBBM/WUS2. Images were taken 13 days after transformation (DAT), 29 DAT, 43 DAT, and 127 DAT for mature plants. Representative cobs from reciprocal crosses of T0 plants (female, pollen acceptor) were pollinated with wild-type (WT) B104 pollen (male, pollen donor), and WT B104 plants (female, pollen acceptor) were pollinated with pollen from T0 plants (male, pollen donor). The number of immature embryos forming RUBY-expressing morphogenic structures relative to the number transformed is indicated and coloured in shades of green according to the percentage. Scale bars: 13 DAT 1 mm, 29 and 43 DAT 1 cm.

In conclusion, of the 60 selected genes, five candidates from four different TF families (AP2/EREBP, GRF, bHLH, homeodomain-leucine zipper (HD-Zip)) consistently and significantly induced RUBY-expressing morphogenic structures reminiscent of those induced by ZmBBM/WUS.

We continued to follow the tissue culture of a subset of candidates that appeared promising in the primary screen until resulting transgenic plants reached full maturity. At 29 DAT, we could observe the emergence of one or more RUBY-expressing shoots emerging from some explants (Fig. 4). We also observed RUBY-negative regenerating plantlets. As we are using phosphinotricin as a selection agent, two out of three regenerants are indeed expected to be an escape (Aesaert *et al*., 2022). In our positive control ZmBBM/WUS2, explants developed large, betalain-accumulating calli and none of the explants developed transgenic shoots at this stage. This ZmBBM/WUS2 phenotype is well-known and a reason why MRs are often used in combination with tissue-specific expression and/or a Cre/loxP auto-excision system (Lowe *et al*., 2016). For most candidates, the rapid emergence of morphogenic structures translated into an increase in RUBY-expressing shoots compared to the tdT control, albeit not significant in this small scale setup. This was the case for bHLH48, EREB152, GRF4, GRF5, HB77, MYB31 (Zm00001eb103730), WRKY53 (Zm00001eb312870) (Fig. 4, Supplementary Table 5). Candidate EREB152, related to ZmBBM (Extended Data Fig. 4), also formed large RUBY-expressing calli, but with one in four explants forming transgenic shoots. Next, we transferred a limited number of plantlets displaying both RUBY-positive shoots and roots to soil in order to detect any possible pleiotropic developmental effects due to the ectopic expression. We found that EREB152, GRF4, GRF5, HB77, and tdT had normal development, while plants expressing bHLH48, MYB31, and WRKY53 displayed mild abnormalities such as stunted growth. Nevertheless, plants were fertile, with the exception of MYB31, that remained stunted, and one plant from both HB77 and bHLH48 (Fig. 4). The ability to obtain seeds from all crosses suggests that continuous overexpression of the candidates does not impact fertility, which is advantageous for gene editing applications, as it enables the segregation of transgenes in subsequent generations after editing (Vandeputte *et al*., 2024). In conclusion, for six TFs (bHLH48, EREB152, GRF4, GRF5, HB77, WRKY53) fertile plants were obtained in the greenhouse.

### Identification of bHLH48, EREB152, GRF4, and HB77 as morphogenic genes to improve maize transformation

Constitutive expression of MRs can lead to pleiotropic developmental effects in developing plants, but also hamper regeneration as observed for ZmBBM/WUS2 and EREB152. Therefore, for further validation and development of these TFs for practical use in maize transformation, we included a Cre/loxP auto-excision system in vector pG3K-Cre-AG-RUBY (Fig. 5A). G418/nptII-selection has been shown to better eliminate escapes (Fig. 5A, Fontanet-Manzaneque *et al*., 2024; Lee *et al*., 2023). In addition, we increased the explant count from 20 to 200, utilizing embryos from three different cobs, to facilitate improved statistical analysis of the transformation frequency. We also focused on the four most promising candidates: bHLH48, EREB152, GRF4, and HB77 that already showed a reproducibly significant increase in the emergence of morphogenic structures.

**Figure 5.**
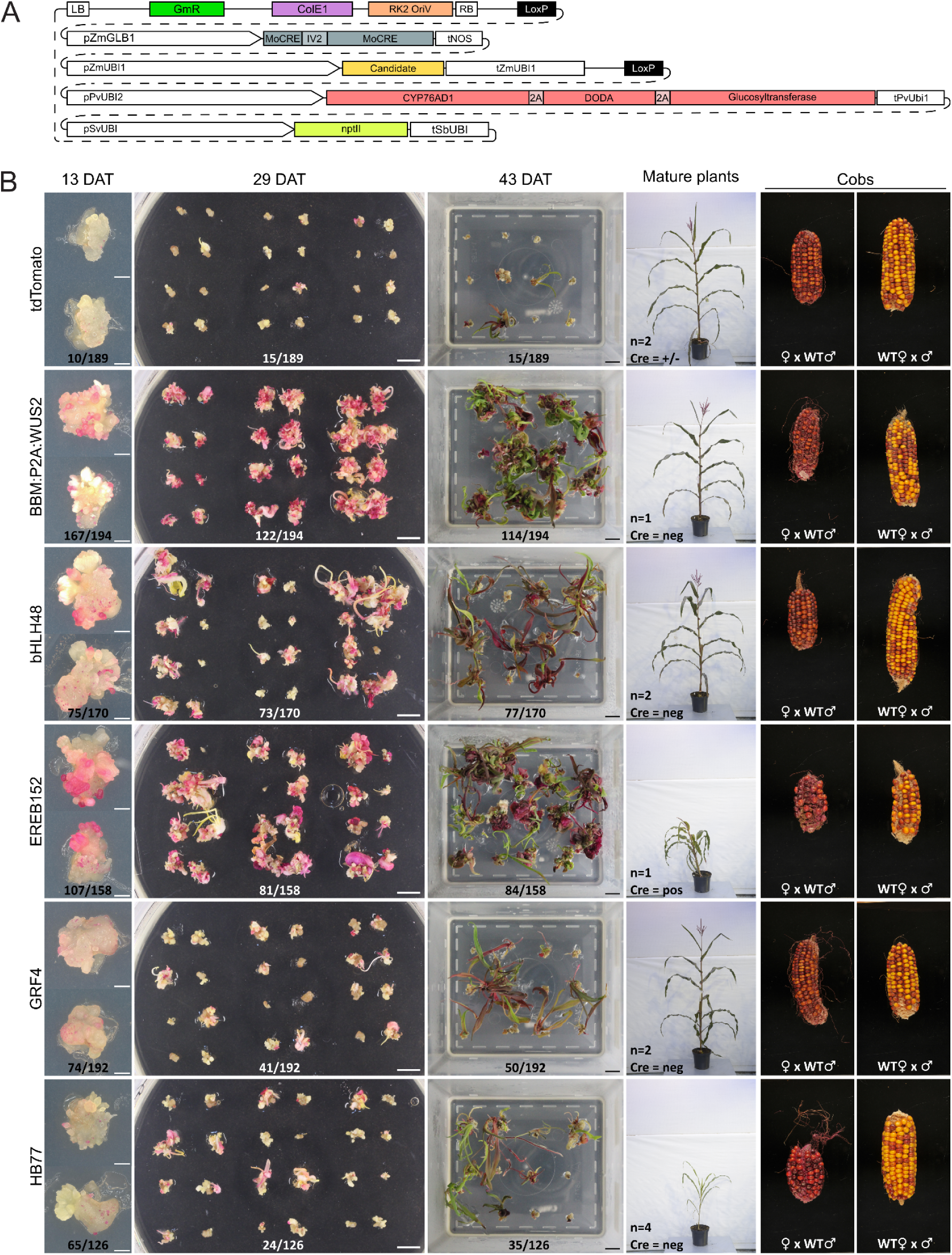
Validation of morphogenic regulators for maize transformation. **A**, Diagram of pG3K-Cre-AG-RUBY. GmR, Gentamicin resistance; ColE1 and RK2 OriV, origin of replication sequences; RB, right border; pZmGLB1, maize *GLOBULIN-1* promoter; MoCRE, Monocot codon-optimized Cre recombinase with intron IV2 of the potato gene ST-LS1; tNOS, *Agrobacterium tumefaciens* nopaline synthase terminator; pZmUBI1 and tZmUBI1, maize UBIQUITIN promoter and terminator; pPvUBI2 and tPvUBI1, *Panicum virgatum L.* UBIQUITIN promoter and terminator; CYP76AD1, P450 CYP76AD1; DODA, l-DOPA 4,5-dioxygenase; pSvUBI and tSbUBI, *Setaria viridis* UBIQUITIN promoter and *Sorghum bicolor* UBIQUITIN terminator; nptII, neomycin phosphotransferase II; LB, left border. **B**, B104 immature embryos were transformed using the Cre/loxP vector equipped with the coding sequence of one of four candidate transcription factors under control of pZmUBI. Shown are representative observed morphogenic induced structures 13 days after transformation (DAT), 29 DAT, and 43 DAT. For tdT, explants from cobs 1, 2 and 3 are shown for reference at 29 DAT. Mature plants of selected high quality events with a single T-DNA copy were imaged at 127 DAT with the MR excised based on Cre being present (pos), absent (neg) or partial excision (+/-). Cobs, reciprocal crosses of T0 plants (female, pollen acceptor) pollinated with wild-type (WT) B104 pollen (male, pollen donor), and WT B104 plants (female, pollen acceptor) pollinated with pollen from T0 plants (male, pollen donor). The positive control construct BBM:P2A:WUS2, which consists of ZmBBM fused to ZmWUS2 via a P2A ribosomal skipping sequence, and the negative control Tandem dimer Tomato (tdTomato) are both driven by pZmUBI. 13 DAT the number of immature embryos forming RUBY-expressing morphogenic structures relative to the amount transformed is indicated. 29 and 43 DAT the number of immature embryos forming RUBY-expressing shoot(s) relative to the amount transformed is indicated. Scale bars: 13 DAT 1 mm, 29 and 43 DAT 1 cm.

At 13 DAT, all four candidates again showed a significant increase in RUBY-expressing morphogenic structures with our new constructs (Fig. 5B). Compared to on average 10 out of 189 explants (5%) for the tdT control, we observed embryo-like structures in 44% for bHLH48, 68% for EREB152, 39% for GRF4, and 52% for HB77, compared to 86% for the ZmBBM/WUS2 positive control (Supplementary Table 6). At 43 DAT, this translated to a much higher transformation frequency compared to 8% for the tdT negative control, significantly increasing three-fold (HB77 and GRF4), five-fold (bHLH48), six-fold (EREB152), and sevenfold for the ZmBBM/WUS2 positive control (Fig. 5B, Supplementary Table 7). As each explant can regenerate multiple RUBY-positive shoots, the transformation frequencies underestimate useful events for gene editing as we have examined previously (Vandeputte *et al*., 2024). Ten plantlets per candidate were pre-selected based on moderate RUBY expression levels, which can be indicative of a single-copy T-DNA. Next, we used digital PCR targeting *nptII* to identify four single T-DNA copy plants for HB77 and ZmBBM/WUS2, three for candidates GFR4 and EREB152, and two for bHLH48 and tdT. By checking presence of MoCre, we could confirm excision of the MR in two plants for HB77, GFR4, and ZmBBM/WUS2 and one for bHLH48 and tdT (Supplementary Table 8). These plants were subsequently transferred to the greenhouse for further growth until maturity and backcrossed to wild-type. In conclusion, we could validate the practical use of bHLH48, EREB152, GRF4, and HB77 as tools to increase maize transformation frequency and generate high quality events.

## Discussion

Here we report on a strategy to uncover regulators of somatic embryogenesis. After genetic transformation of explants with ZmBBM/WUS, SEs are rapidly induced and individual SE cells were isolated using FACS for subsequent single-cell partitioning and mRNA sequencing. We show that prioritization of clusters by automated text mining and ranking of master regulators using MINI-EX, allowed us to make a shortlist of 60 candidate transcription factors for functional analysis. While similar approaches have been used to study human induced pluripotent stem cells at the single-cell levels (Cuomo *et al*., 2020), study of plant somatic embryogenesis has relied on genetics (McFarland *et al*., 2023), bulk RNA-seq (Salvo *et al*., 2014), and/or computational approaches utilizing publicly available datasets (Jiang *et al*., 2024). The latter study predicted 50 maize MRs based on a literature review and multi-omics data from other plant species. Of these MRs, 32 were also identified as regulators in our analysis. Among them, four (DOF11 (Zm00001eb029220), DOF22 (Zm00001eb250390), EREB152, and NAC116) were selected as candidate regulators of somatic embryogenesis, with EREB152 exhibiting a significant phenotype in our screen. This demonstrates that our approach complements existing methods. Our single-cell strategy allows us to achieve higher granularity and enable the identification of cell types of interest. For example, we were able to detect endogenous *ZmBBM* as differentially expressed in somatic embryogenesis-related clusters, which was not observed in a bulk RNA-seq approach (Salvo *et al*., 2014). Furthermore, MINI-EX enabled the prioritization of TFs that were expressed, but not necessarily differentially expressed, and predicted to regulate genes in the clusters of interest. For example, *GRF5* was not differentially expressed in any somatic embryogenesis-related clusters, and three genes *HB77*, *MYB31*, and *WRKY53* were only differentially expressed in one cluster of interest, despite being expressed in several others. Overall, our strategy proved highly successful and we anticipate that this approach can be applied to other MRs, species, and relevant explants for plant transformation, such as leaf fragments (Wang *et al*., 2023). In this way it may help identify MRs adapted to other experimental conditions. In the future, several aspects could be refined to improve its accuracy and applicability of our strategy. For example, ZmWUS2 was not identified as a regulator by MINI-EX because its interactions with target genes lacked support from the available motif information, suggesting that other potential MRs may have been overlooked. This issue can be addressed with improvement of TF motif databases. Moreover, as automated text mining was the primary source for prioritizing relevant single-cell clusters and the candidates within them, further improvements in this technique, such as applying better filters to eliminate false positives or incorporating potentially missing or complementary information could improve the prioritization of MRs (De Clercq *et al*., 2021).

Unsupervised MINI-EX analysis independently pinpointed *ZmBBM* as a morphogenic gene, which was encouraging. HB77 is a HD-Zip factor, and no ortholog has yet been linked to somatic embryogenesis. In contrast, *HB77* has recently been associated with drought stress, and maize *hb77* knock-out lines have a reduction in the number of seminal roots (Yu *et al*., 2024). GRF proteins have been reported as morphogenic genes in wheat (Debernardi *et al*., 2021) and ZmGRF5-LIKE (ZmGRF10, Zm00001eb193180) in maize (Kong *et al*., 2022). Recently, we have shown that ZmGRF1 (Zm00001eb251820) as a chimera with its regulator GIF1 also boosts maize transformation (Vandeputte *et al*., 2024). It is speculated that GRF proteins act in a different pathway compared to ZmBBM and can be combined (Chen *et al*., 2022). bHLH48 is part of Clade Vb (Heim *et al*., 2003) and no related bHLH genes have been shown to act in somatic embryogenesis. Nevertheless, we here consider bHLH48 as the most interesting MR for further development. It combines a strong tissue culture response with a 5-fold increase in transformation frequency, and no signs of pleiotropic effects. Although we only continued with four candidates for in-depth validation, we anticipate that others such as MYB31 and WRKY53 may also have potential as MRs. Indeed, the identification of these MRs is just the first step in developing them as practical tools for crop transformation and gene editing. As with ZmBBM/WUS2 or GRF-GIF, further research is needed to optimize transcriptional units, combine them with other MRs, and test functionality in other genotypes crop species and explants. We anticipate that the expansion in the range of available MRs to enhance plant transformation will play a key role in making plant biotechnology more accessible.

## Methods

### Vector construction and gene synthesis

The plasmids pG3R-WOY-SI, pG3B-SI-AG-RUBY and pG3K-Cre-AG-RUBY were constructed by a combination of Golden Gate cloning and Gibson assembly (Vandeputte *et al*., 2024). Briefly, required DNA elements are assembled into entry vectors with the GreenGate (Lampropoulos 2013) unique overhangs and ligated into the Gibson entry vectors with Unique Nucleotide Sequences (UNS). The Gibson entry vectors are digested with I-SceI and through Gibson assembly with the UNS overlap assembled in the destination vector pG3-U1-AG-U9 (Vandeputte *et al*., 2024). Within these destination vectors, a candidate slot was prepared holding a superfolder GFP (sfGFP) expressed by the *Escherichia coli* glpT promoter and flanked by Golden Gate overhangs, allowing easy identification of clones holding our gene of interest. All used plasmids are listed in Supplementary Table 2. The coding sequences of all candidate genes used were obtained from the Maize genome database (Zm-B73-REFERENCE-NAM-5.0) and prepared for gene synthesis (Twist Bioscience). Only for Zm00001eb034700 a genomic sequence was used, with the intron retained. If required, codon optimization was performed to remove BsaI and AarI restriction enzyme recognition sites and/or to remove specific hard to synthesize regions, such as high G/C content and repetitive elements, while ensuring that the amino acid sequence remained unaltered. Golden Gate C- and D-overhangs and BsaI recognition sites were included during synthesis. All fragments were cloned in a pTwist Amp high copy vector and were directly used as Golden Gate entry clones. List of the sequences of all genetic elements synthesized is found in Supplementary Table 9.

### Plasmid Assembly and Validation in E. coli and Agrobacterium

Constructs were transformed via heat shock in *E. coli* strain DH5α (Invitrogen, Carlsbad, USA) and DH10B (NEB, Ipswich, USA) for larger constructs. We validated the constructs using a combination of restriction enzyme digestion, Sanger sequencing (Eurofins, Ebersberg, Germany), and whole plasmid sequencing (Eurofins, Ebersberg, Germany). Upon construction, expression plasmids were transformed into hypervirulent disarmed *Agrobacterium tumefaciens* strain EHA105 recA^−^ pVS1-VIR2 (Vandeputte *et al*., 2024). The expression vector was isolated from *Agrobacterium* and again validated through whole plasmid sequencing to ensure identity and stability.

### Plant material and growth

Maize growth was performed as described (Aesaert *et al*., 2022). Briefly, *Zea mays L.* (maize) inbred wildtype line B104 seeds (accession no. PI 594047) were germinated in pre-wetted Jiffy-7® pellets under controlled conditions (300 μmol m² s⁻¹, 16 h light/8 h dark, 23°C/22°C). After three weeks, the seedlings were transplanted into medium-sized pots containing professional potting mix (Van Israel nv). After another three weeks, they were transferred to 10L pots and supplemented with controlled release fertilizer (2.0 kg/m³, Osmocote®; Scotts International B.V., Heerlen, the Netherlands, NPK 12/14/24) and grown to maturity in a glasshouse over a period of 70 days (16 h light/8 h dark, 25°C/22°C). To prevent cross-pollination, the ears were covered with paper bags. One day prior to pollination, the silks were trimmed (3–5 cm from the top). On the day of pollination, pollen was collected in a paper tassel bag and applied to the trimmed silks by gently shaking the bag.

### Maize transformation

Maize transformation was performed as described by Vandeputte and colleagues (2024). Briefly, wild-type inbred B104 maize plants were grown in a greenhouse and pollinated. Ears were removed 11-12 days after pollination and stored in a refrigerator until the next morning for *Agrobacterium*-mediated transformation. Furthermore, on the day prior to transformation, the *Agrobacterium* EHA105 recA^−^ pVS1-VIR2 strains holding the expression vectors were cultured on YP plates supplemented with rifampicin, gentamicin, and spectinomycin and were transferred on fresh YP plates with antibiotics. On the transformation day, a loop was taken from the fresh plates and placed in Chu N6-infection medium for 2-5 hours. Immature zygotic embryos derived from at least three different ears were isolated and co-cultivated with the *Agrobacterium* strains carrying the candidate MR genes. Three days after transformation (DAT), zygotic embryos were transferred to resting media supplemented with cefotaxime and vancomycin (antibiotics effective against *Agrobacterium*). At DAT 9, embryos were transferred to a selection medium containing herbicide (phosphinothricin or imazapyr) or antibiotic (geneticin), depending on the selection marker used. At DAT 16, embryos were moved to maturation I medium to further promote development into plants. Surviving embryos at DAT 30 were placed on maturation II medium. Finally, at DAT 44 they were transferred to regeneration II medium. All media are as described (Fontanet-Monzaneque *et al*., 2024).

### Protoplast isolation and single-cell RNA sequencing

Protoplast isolation from transformed zygotic embryos was performed seven days after transformation. We screened the initial 300 zygotic embryos using the Leica M165 FC Fluorescent Stereo Microscope and selected the 200 zygotic embryos expressing the highest fluorescence of yellow fluorescent protein (YFP). The enzyme solution comprised 0.65 M mannitol, 10 mM KCl, 10 mM MES (pH 5.7), 1.25% (wt/vol) cellulase RS, 0.5 % pectolyase Y23, 0.5% macerozyme R-10, adjusted to pH 5.7. Enzymes were activated at 50°C for 10 minutes. After cooling to room temperature, 0.1% bovine serum albumin (BSA), 0.1% CaCl_2_, and 0.009% β-mercaptoethanol were added. Each zygotic embryo was sliced approximately twenty times using a razor blade and incubated in enzyme solution for 60 minutes at 29°C whilst shaking at 40 rpm in the dark. The protoplasts were filtered through a 40 µm filter and washed with the washing buffer containing 0.65 M mannitol and 10 mM MOPS (pH 5.7). Viability was assessed using propidium iodide (PI). Protoplasts without PI signal and positive for YFP were sorted using a BD FACSDiscover™ S8 Cell Sorter for further analysis.

Single-cell RNA sequencing (scRNA-seq) was performed by the Single Cell core facility at VIB according to the 10x Genomics guidelines. Briefly, sorted protoplasts were centrifuged at 400g at 4 °C and resuspended to a concentration of 1000 cells per µL. The cell suspension was processed using the 10x Chromium NextGEM Single Cell 3′ Reagent Kit (V3.1 chemistry, 10X Genomics). Sequencing libraries were prepared and sequenced on an Illumina NovaSeq6000 platform at the VIB Nucleomics Core (VIB, Leuven). An overview of the composition and generation of the scRNA-seq data can be found in Supplementary Table 10.

### scRNA-seq filtering and processing

The raw reads were mapped to the Zm-B73-REFERENCE-NAM-5.0 genome using 10x Genomics Cell Ranger v6.0.0. The resulting gene-to-cell matrix was further filtered by removing genes expressed in fewer than 3 cells; cells with a gene count outside the range 900 to 9,000; cells with library sizes outside the range 1,500 to 60,000; and cells with mitochondrial gene proportions exceeding 25% or chloroplast gene proportions exceeding 15%. Among the remaining cells, only those with transcripts of at least one of the transgenes (*ZmBBM*, *ZmWUS2*, *GUS*, *YFP,* or *HRA*) were retained. The filtered gene-to-cell matrix was further processed using Seurat v4.1.0 (Hao *et al*., 2021). To reduce clustering based on cell cycle-related genes, cell cycle regression was performed using Seurat’s SCTransform function with the argument ‘return.only.var.genes=F’. PCA dimensionality reduction was conducted using Seurat’s RunPCA with ‘npcs=60’. The k-nearest neighbors graph was constructed using Seurat’s FindNeighbors with ‘dims=1:30’. Clusters were identified using Seurat’s FindClusters with ‘resolution=0.8’ and visualized using Seurat’s RunUMAP with ‘dims=1:30’. Cells identified by Cell Ranger as background noise formed an isolated cluster and were removed, after which all previous steps were repeated on the remaining cells. Cluster identities were assigned to the 17 identified clusters by evaluating the expression of known cell-type markers (Supplementary Table 1A).

### Text mining

Known species-gene-trait triples were retrieved from publicly available literature using gene and species annotations identified by PubTator Central (Wei *et al*., 2019). The complete set of PubTator annotations was downloaded on April 11, 2023. The following sections of each downloaded article were processed: title, abstract, introduction, results, discussion, conclusion, materials and methods, and supplementary material. In articles containing relevant species and gene annotations, plant traits were identified using spaCy v3.7.4 (Honnibal *et al*., 2020) with the ‘en_core_web_sm’ trained pipeline. A PhraseMatcher was constructed with the attribute ‘attr=”LEMMA”’ using plant trait synonyms retrieved from the following ontologies: Gene Ontology (Ashburner *et al*., 2000; The Gene Ontology Consortium, 2023), restricted to biological processes only, Plant Trait Ontology (Cooper *et al*., 2018), and Plant Phenotype and Trait Ontology (Liu *et al*., 2023). The identified matches were filtered to only keep those classified as “NOUN” or “PROPN”. A species-gene-trait triple, hereafter referred to as ‘triple’, was created when an occurrence of a plant species, a gene associated with that species, and a plant trait was found within the same sentence, the same paragraph, or if the species name was found in the document title. One triple can be found multiple times within one or several documents, and is characterized by its total number of occurrences.

From the total set of collected triples, only those related to somatic embryogenesis were selected. The full list of selected somatic embryogenesis-related traits can be found in Supplementary Table 1B. In addition to triples collected for maize, triples collected for the model species *Arabidopsis thaliana* were also included, by translating their gene identifiers to those of maize through gene orthology. Orthology information was downloaded from PLAZA Dicots 5.0 (Van Bel *et al*., 2022). One *Arabidopsis* gene was considered as orthologous to one maize gene if the orthology was confirmed by at least two orthologous methods (tree-based, orthologous gene family or best hits and inparalogs). Two sets of orthologous triples were created: one-to-one, consisting of orthologous links connecting exactly one *Arabidopsis* gene to one maize gene, and one-to-many, connecting one *Arabidopsis* gene to multiple maize genes.

To evaluate the quality of collected triples, 35 randomly selected maize triples associated with somatic embryogenesis were manually verified. For each triple, the original text and total number of occurrences were examined. This led to the identification of three criteria: 1) removal of triples found only in Materials and Methods sections, 2) a minimum of 10 occurrences for maize triples, and 3) a minimum of 20 occurrences for *Arabidopsis* triples, increasing accuracy from 51% to 81%.

These filters were applied to generate three gene sets, referred to as priors: 1) P1, composed of triples found for maize; 2) P2, composed of triples found for *Arabidopsis* and translated to maize using one-to-one orthology links, *i.e.*, where one *Arabidopsis* gene corresponds to exactly one maize gene; and 3) P3, composed of triples found for *Arabidopsis* and translated to maize using one-to-many orthology links, where one *Arabidopsis* gene can have one or multiple maize orthologs. Additionally, for further gene regulatory network analysis and gene prioritization, up-regulated genes from (Salvo *et al*., 2014) were used as prior P0. Four additional priors were created by combining priors P0 to P3: P4 was the intersection of P0 and P3, P5 was the union of P1, P2 and P4, P6 was the union of P1 and P2, and P7 was the union of P1 and P3 (Fig. 2B). A list of 38 manually verified genes (Supplementary Table 1C) from the P1 dataset was added to each prior.

### Inference of cell-type-specific regulons

The cell-type-specific regulons for the somatic embryogenesis single-cell dataset were inferred and ranked using MINI-EX v2.0 (Ferrari *et al*., 2022; Staut *et al*., 2023). The filtered Seurat object was used to extract the input data needed for MINI-EX by following the ‘Prepare your files’ document available on the MINI-EX GitHub page. Eight gene-trait files, corresponding to the eight priors, were prepared for prioritization by MINI-EX, by setting the first column to the unique value ‘GO:0010262’ and the last column to the unique value ‘somatic embryogenesis’. The unique term of interest for all the MINI-EX runs was ‘somatic embryogenesis’. The remaining arguments of MINI-EX were used with their default values. MINI-EX was run separately on each prior.

The performance of MINI-EX on the eight priors was evaluated using a gold standard of TFs known to be involved in somatic embryogenesis. This gold standard was constructed by collecting somatic embryogenesis-related TFs that were confirmed by distinct sources: either by text mining independently for maize and *Arabidopsis*, or by text mining and the bulk experiment of Salvo and collaborators (2014). The resulting list was then manually verified to keep a total of 36 gold standard TFs (Supplementary Table 1D).

The clusters most associated with somatic embryogenesis were identified by performing an enrichment analysis (hypergeometric test, FDR-corrected p < 0.05) of the top 100 up-regulated differentially expressed genes in each single-cell cluster against the collected gene sets and somatic embryogenesis-related TF families (Fig. 3B).

### Candidate selection criteria

From the genes obtained, we applied filters to increase transformation potential. First, genes larger than 1500 bp were excluded due to cost considerations for synthesis. Second, each candidate gene underwent a literature review and intellectual property search, supplemented by data obtained from text mining, to discover potential existing characterization. Finally, orthology with *Arabidopsis* and rice (*Oryza sativa subsp. japonica*) was assessed to limit the risk of selecting pseudogenes or genomic annotation errors.

### Microscopy

Transformed embryos were examined for expression of fluorescent protein (YFP for scRNA-seq and tdTomato for negative control) using the Leica M165 FC Fluorescent Stereo Microscope. Images of the embryos at 10, 13, 15, 17 DAT were made using the digital Keyence VHX-7000 microscope using the stitching feature. Images at 22 and 29 DAT were made using a Canon EOS 60D. Finally, images of 43 DAT and maturate plant stages were photographed by a Canon EOS 1200D and/or Canon EOS M50 Mark II.

### Digital PCR

Nanoplate-based digital PCR (dPCR) was used to determine the copy number and presence of backbone. Genomic DNA (gDNA) was isolated from maize seedlings using the Wizard® Genomic DNA Purification kit (Promega), quantified using the Qubit dsDNA HS Assay Kit (Thermo), diluted to 5 ng/μl and added together with primers (8 µM each) and probes (4 µM) to the QIAcuity Probe PCR Kit Master Mix (Qiagen), and subsequently digested with 5 units CviQI (NEB) for 10 minutes at 24°C. dPCR analysis was performed using 8,5K 96-well Nanoplate on the Qiagen QIAcuity One 5-plex platform. PrimeTime™ Probes (IDT, Leuven, Belgium) were equipped with 6-FAM, HEX, and Cy5 fluorophores for the selection marker (*PAT* or nptII), the reference gene *FPGS* (Zm00007a00000670), and the backbone marker (GmR), respectively. All primers and probes used were as described (Fontanet-Monzaneque *et al*., 2024).

## Data availability

Raw and processed scRNA-seq sequencing data will be submitted to GEO and accessible via Plant sc-Atlas (https://bioit3.irc.ugent.be/plant-sc-atlas/) upon submission. MINI-EX and text mining-related files will be posted on Zenodo upon submission.

## Code availability

Code will be made available upon submission.

## List of Figures

**Extended Data Fig. 1.** Dotplot visualizing marker expression per cluster.

**Extended Data Fig. 2.** Expression of bHLH family members results in a similar morphogenic response.

**Extended Data Fig. 3.** Detailed images of morphogenic responses.

**Extended Data Fig. 4.** Phylogenetic trees showing relationship of morphogenic regulators and closest homologs from various species.

## List of Tables

**Supplementary Table 1.** Overview of data analysis

**Supplementary Table 2.** Plasmids used in this study

**Supplementary Table 3.** Induction of morphogenic structures in primary screen

**Supplementary Table 4.** Induction of morphogenic structures in confirmation screen

**Supplementary Table 5.** Preliminary transformation frequency from screen

**Supplementary Table 6.** Induction of morphogenic structures in validation experiment

**Supplementary Table 7.** Transformation frequency in validation experiment

**Supplementary Table 8.** Cre excision in single T-DNA copy plants

**Supplementary Table 9.** List of sequences of synthesized candidate genes

**Supplementary Table 10.** scRNA-seq samples and metrics

## Acknowledgements

We thank Gert Van Isterdael of the VIB Flow Core Ghent for advice on FACS sorting, Karel Spruyt for photography, and Wout Vandeputte for discussions. We are thankful to the VIB Flow Core Ghent, VIB Single Cell Core, and VIB Nucleomics Core for support and access to the instrument park (vib.be/technologies). This work was supported by an Industrieel Onderzoeksfonds grant from Ghent University (F2020/IOF-StarTT/151; IOF.PRO.2021.0017) to S.L. and K.V.

## Author contributions

L.P. and K.V. designed and supervised the research. R.B-A, C.B. designed research. B.D.R. and L.H. supervised research. J.R. performed data analysis, supervised and conducted the cloning and transformation experiments. S.L. performed data analysis. I.V. performed cloning and transformation experiments. S.A., G.C. performed transformation experiments. J.D.B., C.G., and T.E. performed protoplasting, single cell partitioning, and sequencing experiments. All authors discussed the results and commented on the manuscript.

